# One enzyme many faces: alkaline phosphatase-based phosphorus-nutrient strategies and the regulatory cascade revealed by CRISPR/Cas9 gene knockout

**DOI:** 10.1101/2020.05.20.107318

**Authors:** Kaidian Zhang, Zhi Zhou, Jierui Wang, Jiashun Li, Xin Lin, Ling Li, Xiaomei Wu, Yanchun You, Senjie Lin

## Abstract

Phosphorus (P) is an essential macronutrient for marine phytoplankton responsible for ∼50% of global carbon fixation. As P availability is variable and likely will decrease in future warming oceans, phytoplankton growth will be constrained by their strategies to scavenge dissolved organophosphate. To enhance our mechanistic understanding of these strategies, here we employ CRISPR/Cas9 to create mutants of alkaline phosphatase (AP) PhoA and PhoD and a putative regulator in the diatom model *Phaeodactylum tricornutum*, coupled with transcriptomic profiling to interrogate their modes of function and P- regulatory network. Results indicate that these two AP isoforms are differentiated in subcellular localization and substrate specialization, and are mutually compensatory and replaceable. Further analyses reveal a regulatory cascade of P scavenging and potential roles of AP in iron and ammonium uptake as well as diverse metabolic pathways. These findings have important implications in how phytoplankton community will respond to future changing microenvironments of global oceans.

## Introduction

Like land plants, marine phytoplankton contribute ∼50% of global carbon fixation from carbon dioxide that provides organic carbon and oxygen to the worldwide biota (1). The growth and productivity of both land plants and phytoplankton rely on phosphorus (P) and other nutrients (2). However, in various parts of the world’s oceans, P-nutrient, primarily dissolved inorganic phosphorus (DIP), is limited (3-5). To maintain intracellular P homeostasis and population growth, phytoplankton have evolved strategies to utilize the more abundant dissolved organic phosphorus (DOP) as an alternative source of P (6). They mainly achieve this via alkaline phosphatase (AP) that hydrolyses phosphomonoesters, which account for ∼75% of total DOP in the ocean, and other enzyme systems that hydrolyse phosphonates, which account for ∼25% of total DOP (5, 7). These environmental characteristics, along with the short generation times of phytoplankton (unicells), present an excellent model to elucidate how AP confers photosynthetic organisms the ability to utilize the highly diverse organophosphate compounds and maintain population growth in P-poor environments.

Diatoms are one of the most dominant primary producers in the global ocean, contributing about 40% of marine primary production. The rapidly growing genomic studies on marine phytoplankton have resulted in the emergence of two diatom model species, *Phaeodactylum tricornutum* and *Thalassiosira pseudonana*, which received enormous research attention. However, how phytoplankton in general or diatoms in particular regulate the expression and function of AP for scavenging different sources of P, maintaining P homeostasis and responding to low-P stress is poorly understood. Here, we use *Phaeodactylum tricornutum* as a model and employ CRISPR/Cas9 to knock out two AP genes (*PhoA* and *PhoD*) and the gene of SPX protein, a P regulator previously known only in plants (8), and perform transcriptome profiling to characterize their modes of functions. The results unveil striking functional differentiation, compensatory expression, and replaceable regulation of the different isoforms of AP, and the SPX-PHR-PSI regulatory cascade of P-nutrition and homeostasis. Besides these novel insights, our network-based functional interrogation shed light on potential functional interactions of P and AP with uptake of of iron and nitrogen as well as various metabolic pathways. These findings have significant ecological implications regarding how phytoplankton taxa may differentially respond to and compete for the potentially increasingly limited P in future oceans.

## Methods

### Algal culture

*Phaeodactylum tricornutum* strain CCAP 1055/1 was obtained from the Culture Collection of Algae and Protozoa (Scottish Marine Institute, UK; https://www.ccap.ac.uk/). Cells were cultured in autoclaved 0.22-µm filtered oceanic seawater collected offshore in the South China Sea, enriched with the full nutrient regime of the f/2 medium without additional silicic acid. The cultures were made axenic by treatment with 1 × KAS compound antibiotics (50 µg ml^-1^ kanamycin, 100 µg ml^-1^ ampicillin and 50 µg ml^-1^ streptomycin). Cultures were incubated at 20°C under a 14:10 light dark cycle with a photon flux of 100 µE m^−2^ S^−1^.

### PAM-target site selection and vector construction

The PhytoCRISP-Ex, a CRISPR target finding tool, was used to design Cas9 target sites (G-N19-NGG) with low/no off-target potential (9) (Table S1). The single guide RNA (sgRNA) adapters targeting PhoA, PhoD and SPX were individually ligated into separate pKS_diaCas9_sgRNA plasmids (Addgene ID: 74923) as described in Nymark *et al*. (10). The recombined pKS_diaCas9_sgRNA plasmids were co-introduced with pAF6 plasmid carrying Zeocin resistance gene (Invitrogen, Thermo Fisher Scientific, Grand Island, New York, USA), with the former to induce mutation and the latter to facilitate selection (11).

### Biolistic transformation

For co-transformation with the dual plasmids, cells were collected from exponentially growing cultures and were then concentrated to 2 × 10^8^ cells ml^-1^ at 3000 g for 5 min. Then 200 µl of the cell suspension was spread on 1.5% agar plates containing 50% seawater supplemented with f/2 medium nutrients without silica. Transformation was performed using a Bio-Rad Biolistic PDS-1000/He Particle Delivery System (Bio-Rad, Hercules, California, USA) as described previously (12). A burst pressure of 1550 psi and a vacuum of 28 Hg were used. Three mg of assembled pKS_diaCas9_sgRNA plasmid and 3 mg of the pAF6 selection plasmid were used for co-transformation. The bombarded cells were incubated on the agar plate under low light (50 µmol photons m^−2^s^−1^) at 20°C. Two days after the transformation, cells were re-suspended in 600 µL sterile 50% seawater. About 200 µl of this suspension was plated onto agar medium containing 50 µg ml^-1^ Zeocin and 1 × KAS compound antibiotics, and the plate was incubated under a 14:10 light dark cycle at a photon flux of 100 µmol photons m^−2^ s^−1^ at 20°C. After three weeks, the colonies growing on the plate were re-steaked on fresh 9 cm 1% agar plates containing 75 µg ml^-1^ Zeocin to isolate pure mutant strains.

### DNA isolation and genotype characterization

Pure colonies from above were transferred to 24-well plates (Bio-Rad), and cells were incubated for 10 days. To lyse the cells, 100 µl of transformed cells were removed to PCR tubes. Tubes were centrifuged for 2 min and then re-suspended in 50 µl TE buffer (10 mM, pH 8.0). The resuspended cells were frozen in liquid nitrogen, then heated immediately in 98°C for 10 min using PCR Thermal Cycler (Bio-Rad). The freezing-heating cycle was repeated for 3 times. After a brief centrifugation to pellet cell debris, the supernatant for each clone was used directly or stocked at 4°C. Three microliters of cell lysates were used as template for PCR analysis using ExTaq DNA polymerase (Takara, Mountain View, California, USA). For checking the presence or absence of Cas9 gene components in the genome, primer pairs were designed in the 3’ region for PCR amplifying the sequences (Table S2). To confirm the disruption (insertions and deletions) of the target genes, the target regions were amplified using specific primers designed flanking the targets (Table S2). The PCR products were separated by electrophoresis on a 1% agarose gel, purified using MiniBEST Agarose Gel DNA Extraction Kit (Takara), and cloned into pMD19-T (Takara). Random clones were picked for Sanger sequencing.

### Comparison of growth between wild type and mutants

The mutant and wild type (WT) cells were first cultured in f/2 medium until they reached the exponential growth phase. Then, the cultures were inoculated into low-Pi (5 µM) f/2 medium and grown until phosphate in the culture was all depleted (< 0.5 µM). The P-limited mutant cells and WT cells were finally inoculated separately into media with different phosphorous nutrient conditions: P+ (36 µM DIP), phytate (PA) (36 µM), glycerol-3-phosphate (G3P) (36 µM), triethyl phosphate (TEP) (36 µM) and P− (DIP < 0.5 µM), each in triplicate. These DOP compounds were all provided by Sigma-Aldrich (St. Louis, Missouri, USA). In each culture, 1 × KAS compound antibiotics were added to avoid bacterial interference. Cell concentration was determined daily using CytoFLEX flow cytometer (Beckman Coulter, Indianapolis, Indiana, USA), which measures chlorophyll fluorescence with the excitation light of 488 nm and emission at 690 nm. Around 2 × 10^7^ cells from each culture were collected every other day by centrifugation at 3000 g, 4°C for 5 min and re-suspended in 1 ml Trizol Reagent (Thermo Fisher Scientific) and stored at −80°C for subsequent RNA extraction. The supernatant recovered was used to measure DIP concentration with the molybdenum blue method (13).

### The measurement of alkaline phosphatase activity

AP activity was determined by adding 50 µl of 20 mM p-nitro-phenylphosphate (p-NPP from Fluka; dissolved in 1 M Tris buffer at pH 8.5) into 1 ml culture sample to yield final concentrations of 1 mM p-NPP and 50 mM Tris at pH 8.5 (14). Each reaction was carried out in a sterile 1.5 ml tube in the dark at 20°C for 2 h. The samples were then centrifuged at 12000 g for 1 min. The supernatant was used for total AP activity measurement at 405 nm on a SpectraMax Paradigm plate reader (Molecular Devices, San Jose, California, USA). For the measurement of extracellular AP activity, the culture was filtered through a 0.22-µm membrane and the filtrate was used to detect AP activity using the same spectrophotometric method as described above.

### RNA extraction, cDNA synthesis and gene expression measurement

Total RNA was extracted using Trizol Reagent coupled with Qiagen RNeasy Mini kit (Qiagen, Germantown, Maryland, USA) following a previously reported protocol (15). The RNA extracts were treated with RQ1 DNase (Promega, Madison, Wisconsin, USA) to remove any genomic DNA contamination. RNA concentrations were measured using NanoDrop ND-2000 Spectrophotometer. For each sample, 200 ng of total RNA was reverse-transcribed in a 20 µl reaction mixture using the ImProm-II reverse transcriptase (Promega) with random hexamer primer according to the manufacturer’s instructions. Each cDNA preparation was diluted 1: 20 with nuclease-free water for further analysis. To assess the differential expression of each target gene under different conditions, RT-qPCR analyses were conducted using specific primers designed for each target gene (Table S3). RT-qPCR was performed with a CFX96 Touch Real-Time PCR detection system (Bio-Rad) with iQ™ SYBR Green Supermix (Bio-Rad) following the manufacturer’s recommendations. Three replicate assays were performed for each RNA sample.

### Phylogenetic Analysis of AP, PT and SPX genes in *P. tricornutum*

To understand the types of APs, PTs, and SPX domain-containing protein genes identified, the phylogenetic trees were inferred for each gene based on the amino acid sequences of the gene of interest and sequences of previously reported known types of the gene obtained from NCBI NR database. The amino acid sequences were aligned using MUSCLE (16) embedded in the software MEGA X (17), and edited manually using MEGA X. Phylogenetic tree reconstructions were performed employing the PhyML-aLRT method (18) with LG, WAG+G+F and LG+G models for AP, PT and SPX respectively and best of NNI & SPR on the Seaview platform (19).

### Construction and deep sequencing of transcriptome libraries

Samples were collected on the third day from mutant and WT cultures, triplicated for each cell type and growth condition, including Wild Type (WT), mutants of PhoA (*m*PhoA), PhoD (*m*PhoD), and SPX (*m*SPX), each under P-replete (P+), P-depleted (P−) conditions, and glycerol-3-phosphate (G3P), totaling 36 samples. RNA was extracted for each sample and was measured using NanoDrop ND-2000 Spectrophotometer as described above, and RNA integrity was verified with Agilent 2100 Bioanalyzer (Agilent Technologies). Thirty-six single-end fragment libraries (50 bp) were constructed and sequenced on the BGISEQ-500 platform according to the manufacturer’s instructions (BGI). Sequencing reads were processed by trimming off adaptors and primers and removing low-quality reads. The good-quality reads retained were mapped to the annotated genome of *P. tricornutum* strain CCAP 1055/1 to identify genes, quantify expression levels, and analyze coordinated expression networks (see below).

### Weighted gene co-expression network analysis

The genomic sequences and genome annotation of *P. tricornutum* strain CCAP 1055/1 were downloaded from Ensembl Genomes archive sites (http://ensemblgenomes.org/). The alignment of our transcriptome reads to the genome was performed using the HISAT2 software (20). StringTie and DESeq2 software were used to estimate transcript abundances and identify differentially expressed genes among different groups (FDR < 0.05, fold change > 2), respectively (21). Based on the mRNA expression level of all genes in transcriptome libraries and alkaline activities of each culture, the WGCNA package was employed to construct global weighted correlation network (22). The network was further analyzed and displayed using the Cytoscape software (23).

## Results

### Gene knockout reveals differential subcellular distributions of APs

We identified eight AP genes from our *P. tricornutum* transcriptomes (SRP214503), including the previously reported *PhoA* (Phatr3_J49678), five putative *PhoD*, one *PhoA*^*aty*^ and one unclassified AP gene (Fig. S1 and Table S4). Most of these genes were significantly up-regulated under P-stress (P−) (Table S4). *PhoA* and the most P-responsive *PhoD* (Phatr3_J45757) were selected for knockout. We designed multiple single-stranded (sg)RNA sequences for each based on available protospacer adjacent motif (PAM) sites and transformed *P. tricornutum* cells with dual plasmids carrying sgRNA-Cas9 and zeocin resistance, respectively (Fig. S2 and Table S5). Both insertions and deletions (indels) occurred at the target site in each gene (Fig. S2).

Knockout mutants *m*PhoA21 and *m*PhoD30 were selected for investigating the differentiation of subcellular functional locality of PhoA and PhoD. In the P− culture, extracellular AP activity and total AP activity in *m*PhoA decreased by 95% and 76%, respectively (Fig. S3), indicating that PhoA is an extracellular enzyme. When averaged over a 10 day experimental period, PhoA accounted for ∼75% of total AP activity (Fig. S4). By contrast, *PhoD* mutation caused no remarkable changes in either extracellular AP or total AP under P− conditions (Fig. S3). Thus, PhoD is an intracellular AP and only contributes a minor fraction (<25%) of total AP.

### Differential functions of APs and switchable pathways of DOP utilization

To further examine the functional relationship between PhoA and PhoD, three different DOP compounds, phytate, glycerol-3-phosphate (G3P), and triethyl phosphate (TEP), were supplied as the sole P source. Mutants and wild-type (WT) diatoms were grown with one of the three DOPs or DIP (P+), or without P (P−).

Without PhoA, *P. tricornutum* utilized phytate or DIP as the sole P source efficiently, but without PhoD, much less growth occurred (Fig. 1a-d and Fig. S5). Because we found no DIP released into the medium (Fig. 1b), apparently, phytate was not hydrolysed extracellularly, but was transported into the cells for use, where PhoD would play a major role (Fig. 1e). Its import might cost energy, explaining the observed <100% growth efficiency. With TEP as the sole P source, we observed neither growth nor DIP release in the WT or mutant cultures (Fig. 1a, b, d), indicating *P. tricornutum* is unable to utilize TEP (Fig. 1e). By contrast, WT with G3P cultures displayed growth comparable to those of WT with DIP (Fig. 1a, d), indicating that *P. tricornutum* utilizes G3P at 100% efficiency. Meanwhile, a significant amount of DIP was released into the medium, while none was detected in the *m*PhoA group cultured with G3P (Fig. 1c). Therefore, G3P utilization involves extracellular hydrolysis by PhoA. Taken together, PhoA and PhoD are specialized for different DOP substrate types (Fig. 1e).

**Fig. 1.**
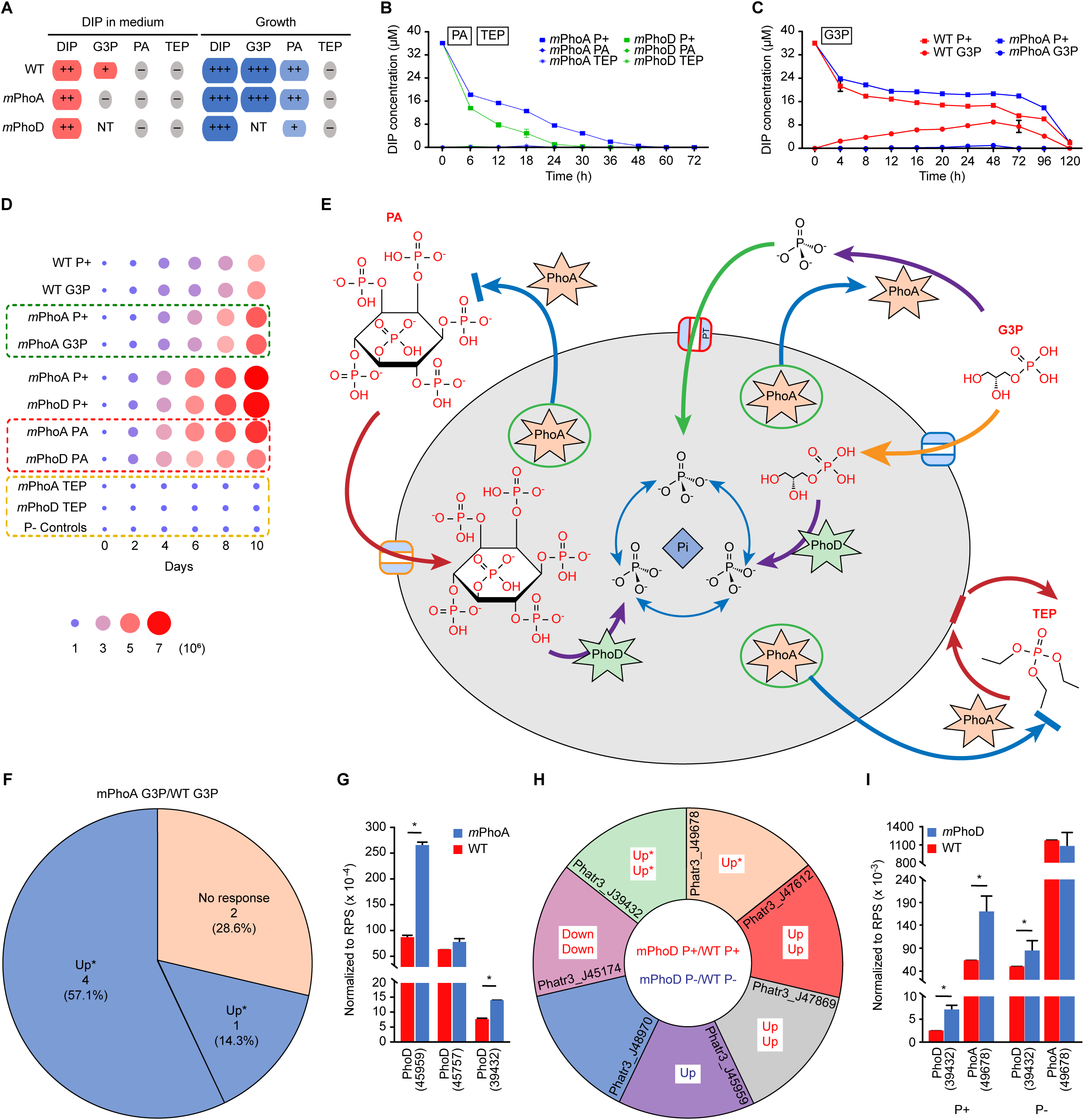
DOP utilization under the coordinated functions of different APs and the compensatory responses between PhoA and PhoD. **a** DIP concentration in the medium and cell growth under different P conditions. Size of the circle and the number of the plus sign corresponds to DIP concentration detected in the medium and the growth rate. The minus sign shows no DIP release in the medium and no cell growth. NT, not tested. **b-c** DIP concentration in the culture. P+, P-replete culture; P−, P-depleted culture; G3P, glycerol-3-phosphate; PA, phytate; TEP, triethyl phosphate. Each data point is the mean of triplicate cultures with the error bar indicating standard deviation. **d** Heatmap of cell growth under different P conditions. The scale on the right represents cell concentration as ×10^6^ cells mL^-1^. P− Controls depicts the low growth of P-deprived mutants and WT cultures. **e** Model of DOP utilization in *P. tricornutum*. G3P can be hydrolysed to release DIP by the secreted PhoA or be imported into cells and utilized via the intracellular PhoD. PA only can be utilized by the intracellular PhoD and TEP cannot be utilized. **f** Relative expression of AP genes in *m*PhoA cultures under G3P condition determined by RNA-seq. **g** Relative expression of AP genes in *m*PhoA under G3P condition determined using RT-qPCR. Asterisk depicts significant difference. **h** Relative expression of AP genes in *m*PhoD determined by RNA-seq. **i** Relative expression of AP genes in *m*PhoD determined by RT-qPCR. Asterisk shows significant difference.

Surprisingly, given the reliance of G3P utilization on PhoA, *PhoA* mutation did not inhibit *P. tricornutum* growth in the G3P medium, but allowed similar growth to that of *m*PhoA with DIP (Fig. 1a, d). This indicates that *P. tricornutum* can switch to a different mechanism when PhoA function is lost. This mechanism was unlikely to be another extracellular hydrolase because no DIP release was detected; therefore, it likely involved importing G3P into the cell and hydrolyzing it via PhoD or other intracellular APs (Fig. 1e).

### Compensatory regulation among AP genes observed from AP mutants

RNA-seq showed that *PhoA* mutation induced up to threefold up-regulation of two PhoD genes and two other putative AP genes under G3P conditions (Fig. 1f and Table S6), a result verified by RT-qPCR (Fig. 1g). Similarly, in *m*PhoD, other AP genes were significantly up-regulated (up to threefold, p<0.05) under P+ or P− conditions, as both RNA-seq and RT-qPCR results showed (Fig. 1h, i and Table S7). Both cases indicate a remarkable mutually compensatory up-regulation between *PhoA* and *PhoD* when one of them is functionally lost.

### SPX mutation uncovers a regulatory cascade of AP and P transporter genes

We identified six genes harboring an SPX domain, which is known as a sensor of intracellular P level and regulator of P homeostasis in land plants (8) but has not been studied in algae; of these, Phatr3_J47434 only contains the SPX domain (SPX protein) and is expected to exclusively regulate P uptake and homeostasis, while others contain additional domains (SPX-like protein) and may regulate a variety of physiologies (8, 24). Our results showed that the SPX and two of the SPX-like genes were inducible by P stress (Fig. S6 and Table S8). We chose to study the SPX gene, and obtained multiple mutants (*m*SPX) with various indels (Fig. S2). *SPX* disruption led to significant increases in AP activities (Fig. 2a, b) and gene expression (Fig. 2c and Table S9) under both P+ and P− conditions. This is reflected in both RNA-seq and RT-qPCR results, with RT-qPCR indicating 3.3−22.4-fold up-regulation for three AP genes in *m*SPX/P+ cultures and 1.4−2.2-fold up-regulation for four AP genes in *m*SPX/P− cultures (Fig. 2d). This is clear evidence that SPX is an upstream negative regulator of AP genes.

**Fig. 2.**
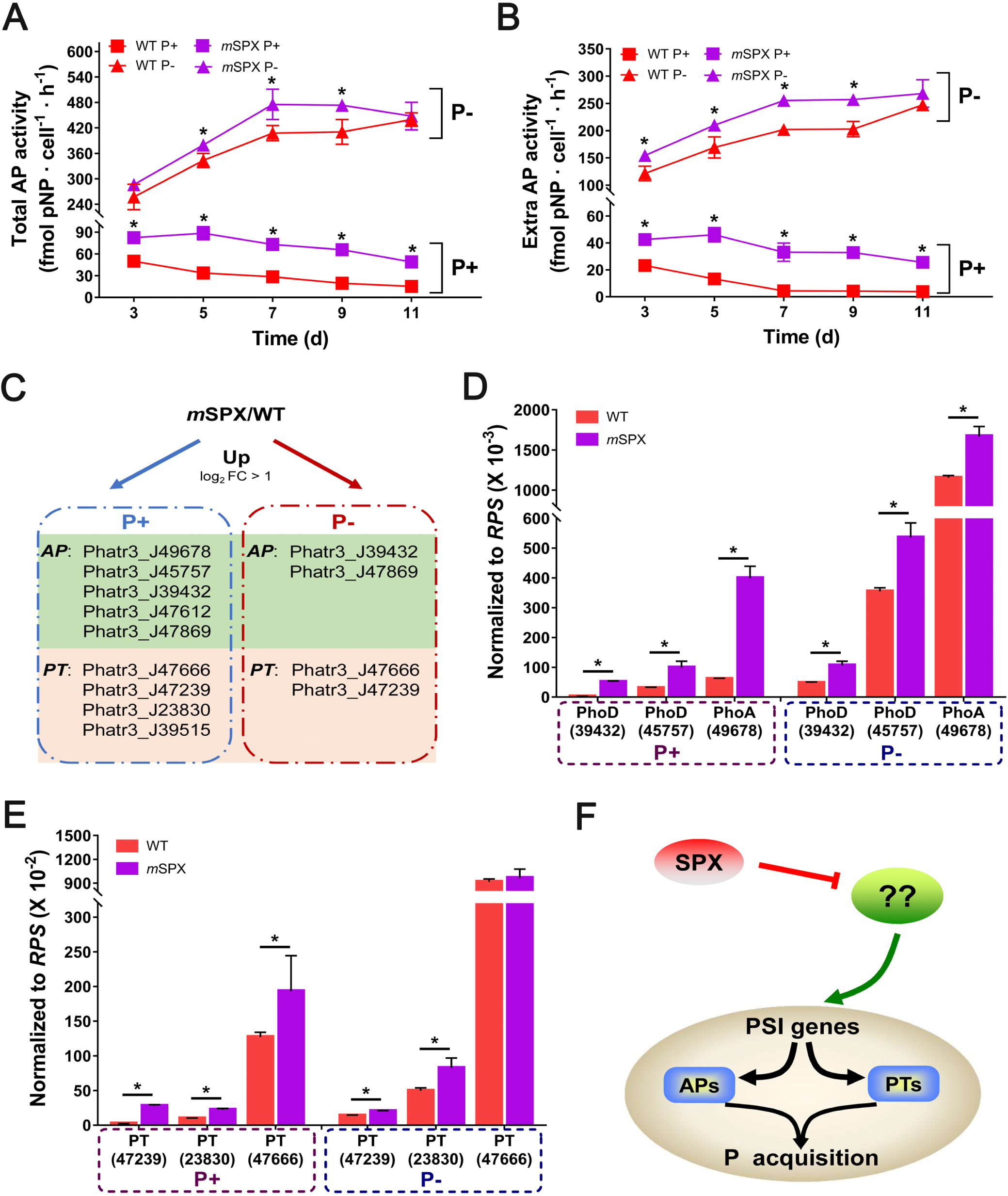
Evidence that SPX is upstream negative regulator of AP and P transporters. **a** Total AP activities in *m*SPX strains. **b** Extracellular AP activities in *m*SPX strains. **c** The significantly up-regulated AP and PT genes determined by RNA-seq in *m*SPX under P+ and P− conditions. **d** The relative expression of AP genes in *m*SPX determined by RT-qPCR. **e** The relative expression PT genes in *m*SPX determined by RT-qPCR. **f** The model of SPX as an upstream negative regulator in *P. tricornutum*. Positive and negative effects are indicated by arrows and flat-ended lines, respectively. Asterisk depicts significant difference.

Furthermore, our results showed that SPX is also a regulator of P transport. We identified 13 phosphate transporter (PT) genes from our transcriptomes, including seven encoding sodium-dependent phosphate transporters (SPT) and six major facilitator superfamily inorganic phosphate transporters (IPT). Under P− conditions, six of the SPT genes showed significant up-regulation, while only one showed no change; by contrast, only one *IPT* showed significant up-regulation (Table S10). *SPX* mutation led to significant up-regulation of most of these P-responsive transporter genes regardless of P conditions, as shown by both transcriptomic (Fig. 2c and Table S11) and RT-qPCR data (Fig. 2e). Our phylogenetic analysis confirms that the non-P-responsive transporters (exclusively IPT) are low-affinity transporters functioning under P-rich conditions, while those promoted by P stress and SPX mutation (mostly SPT) are high-affinity transporters (Fig. S7).

Surprisingly, *SPX* in WT was also up-regulated under P stress (Table S8): unexpected for a negative regulator of AP and PT genes. This suggests that AP and PT are not immediate targets of SPX; rather, an intermediate control mechanism may exist between SPX and AP or PT (Fig. 2f). We identified seven genes encoding P starvation response protein (PHR) (25) that were inducible by P stress, one of which (Phatr3_J47256) showed ∼20-fold up-regulation (Table S12). Furthermore, this *PHR* exhibited significant up-regulation in *m*SPX but showed no response to *PhoA* and *PhoD* mutations, indicating that it functions upstream of APs (Table S13). These results place PHR downstream of SPX and upstream of APs and PT genes.

### Broader metabolic pathways influenced by AP, SPX and P-stress

We sequenced 36 RNA samples from 12 experimental conditions (Table S14) to understand what metabolic pathways are impacted by APs, P condition, and SPX. Data were subjected to weighted gene co-expression network analysis (WGCNA) (Fig. S8). One of the subnetworks composing 1622 differentially expressed genes (Table S15) showed strong linkages, and 82 of these genes exhibited particularly strong influences of mutations of *AP* and *SPX*. Of these, 76 genes displayed strong correlation with SPX,indicating a broad range of effectors of this regulator (Fig. 3a and Table S16).

**Fig. 3.**
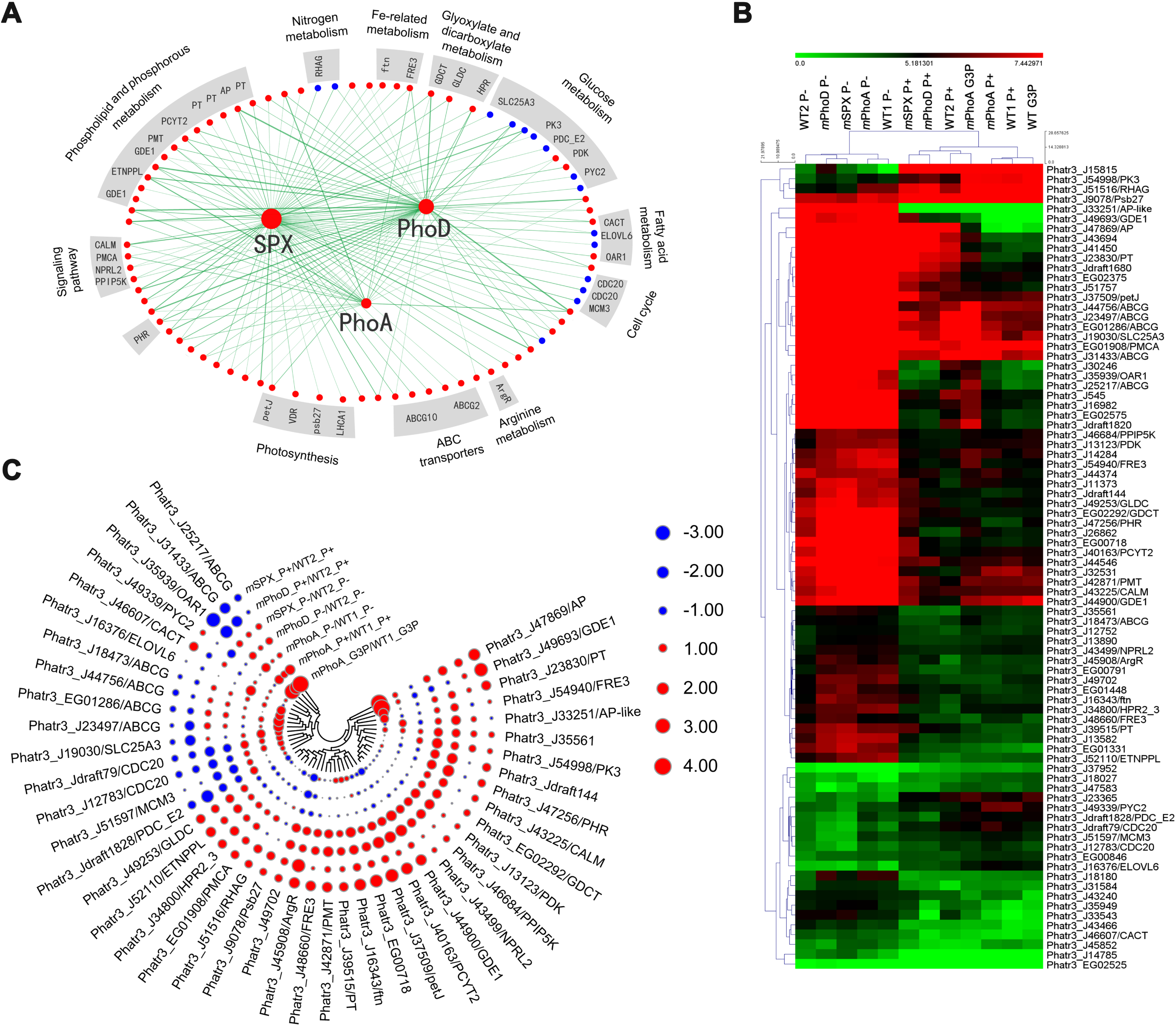
Weight gene co-expression network analysis (WGCNA) and cluster heat map. **a** The network of the 82 most highly connected genes. Several metabolic processes, including phospholipid metabolism and parts of photosynthesis are regulated by P deficiency. The network only displays the first neighbor of the three mutated genes where the corresponding topological overlap is above the threshold of 0.05. Red dots depict positive correlations with AP activity; blue dots depict negative correlations with AP activity. Genes to be elaborated in the main text are shown with abbreviated names. **b** Expression profiles of the 82 core genes classified in the mutant transcriptomes under different P conditions. **c** Comparison between mutant and wild-type groups under different P conditions. The scale on the right represents log2 transformed gene expression fold changes.

Intact AP genes showed strong correlation with mutations of *AP* and *SPX* (Fig. 3a), further demonstrating compensatory regulation among APs. As part of the DOP utilization pathway, several ATP-binding cassette transporters also showed positive correlation with APs and SPX (Fig. 3a), which were remarkably up-regulated under G3P conditions (Fig. 3b). Furthermore, seven phospholipid metabolism-associated genes were up-regulated in *m*PhoD and *m*SPX, in which AP activity increased (mostly due to PhoA) regardless of P presence (Fig. 3b, c and Table S16). These suggest that alteration of the PhoD-PhoA balance and AP-PT regulation via the SPX-PHR cascade modulates phospholipid metabolism, a probable primary responder to P-nutrient dynamics.

The compounded impacts of AP, P, or SPX-dependent factors on metabolic pathways can be disentangled by comparing responses of the metabolic processes to the disruption of these genes and P supply. For example, most of the genes in the glycolysis/gluconeogenesis metabolism subnetwork were negatively correlated with AP activity (Table S16), but either be due to a negative influence of AP or a positive effect of P. However, pyruvate kinase gene, the key enzyme of glycolysis, and pyruvate dehydrogenase E2 component, the rate-limiting enzyme in the TCA cycle, were not repressed under P-deficiency as would be expected if they were under direct control of P (Fig. 3b) but were up-regulated in *m*PhoD/P− and *m*SPX/P− (Fig. 3c). It appeared then, the functional loss of PhoD and SPX was the reason. Given that the mutations of these two genes resulted in up-regulation of intact APs (compensatory regulation shown above) (Fig. 1h, 2c), these results signal direct promoting roles of non-PhoD APs on glycolysis/gluconeogenesis. Even though further experimental verification is needed, such interrogation is meaningful for generating hypotheses.

By the same way of interrogation, our data show that P and APs by acting at different nodes have regulatory influences on various major metabolic pathways. In the photosynthesis pathway, photosystem II Psb27 protein was repressed by P stress (Fig. 3b) and up-regulated in *m*PhoD/P+ and *m*SPX/P+ (Fig. 3c), but showed no change in *m*PhoA, suggesting it is directly regulated by P rather than PhoA. A cytochrome c6 (petJ) was induced by P stress (Fig. 3b) and up-regulated in *m*SPX and *m*PhoD regardless of P condition, but not changed in *m*PhoA, suggesting the involvement of other APs. These results suggest that P and APs acting at different points of the pathway both have regulatory influences on photosynthesis.

A 3-oxoacyl-[acyl-carrier protein] reductase gene important in fatty acid metabolism, showed a positive correlation with P deficiency (Fig. 3b), consistent with the previous finding that fatty acid content increases under P stress (26, 27). It was down-regulated in *m*PhoD/P+ and *m*SPX/P+ groups (Fig. 3c), however, suggestive of a promoting roles of P deficiency in the metabolism. Meanwhile, transcription of a very long chain-fatty acids elongase (ELOVL6) gene was rpressed under P stress (Fig. 3b) but up-regulated in *m*PhoA/P−, evidence that decreases in long-chain fatty acid content under P stress (28) might actually be due to the negative regulation by PhoA. Besides this, a mitochondrial carnitine/acylcarnitine transporter (CACT) was up-regulated in *m*SPX/P+, indicating that SPX can also affect fatty acid oxidation by controlling the transport of carnitine/acylcarnitine (Fig. 3c).

Glyoxylate and dicarboxylate metabolism connects the TCA cycle and various other metabolic pathways (29). Three genes in this metabolism are positively correlated with AP activity (Fig. 3b) and up-regulated in *m*PhoD and *m*SPX, but down-regulated in *m*PhoA, indicating a dominant regulatory role of PhoA in glyoxylate/dicarboxylate metabolism. Similarly, four signaling pathway-related genes (Table S16) are positively correlated with AP activity (Fig. 3b) and showed up-regulation in *m*PhoD and *m*SPX, but no change in *m*PhoA. Thus, this P-related signaling pathway likely responds to P-stress through SPX and APs other than PhoA and PhoD.

We identified nine ammonium transporter genes, four of which were significantly down-regulated under P− conditions (Table S17), indicating P dependency (30) or AP dependency. The up-regulation of RHAG in *m*SPX/P+ and *m*PhoD/P+ relative to WT/P+ (Fig. 3c), however, supports AP (non-PhoD) dependency. Furthermore, an arginine repressor (ArgR) was induced by P stress (Fig. 3b) and up-regulated in *m*PhoD/P+ and *m*SPX/P+, also indicating a positive influence of non-PhoD APs. Evidently, the SPX-PHR-APs (non-PhoD) regulatory cascade influences nitrogen-nutrient and arginine metabolism.

Iron is an essential, and often deficient, micronutrient for oceanic diatoms and other phytoplankton. Interestingly, three iron uptake-related genes (one ferritin and two ferric reductases) showed a positive correlation with P stress (Fig. 3b). Ferritin (ftn), which stores iron and releases it in a controlled fashion, was also up-regulated in *m*PhoD and *m*SPX (Fig. 3c). Similarly, *SPX* mutation promoted ferric reductase genes *FRE3* (Fig. 3c). By contrast, *PhoA* mutation caused no changes in expression of these genes, except under G3P treatment. These results suggest that SPX and non-PhoA non-PhoD APs act in concert in promoting iron uptake under P deficiency. Probably, a cross-talk between P and iron homeostases exists, as does in plants (31, 32), that is mediated by the SPX-PHR-AP mechanism.

Previous studies have shown that P deficiency halts phytoplankton cell division and population growth (33, 34). Accordingly, cell division cycle protein 20, DNA duplication licensing factor MCM3, and co-factor of the anaphase promoting complex in *P. tricornutum* were negatively correlated with AP activity (Fig. 3b). However, their expression was up-regulated in *m*PhoA but down-regulated in *m*PhoD and *m*SPX regardless of P conditions, suggesting a direct role of PhoA.

## Discussion

Although it is well recognized that phytoplankton have evolved a series of mechanisms, such as the AP enzyme system to scavenge DOP (28, 35, 36), how the redundant genes are coordinated and regulated in function to maintain P homeostasis under variable P conditions has remained unclear. Here we document in the model diatom *P. tricornutum* eight AP genes, six SPX genes, seven PHR genes, and thirteen PT genes. Through functional genetic manipulation, transcriptome profiling and physiological observations, we discover APs’ intricate, functional differences and a central regulatory cascade of the P-stress response, depicting their influences on the general metabolic landscape (Fig. 4). These findings have important ecological implications regarding how varying P conditions can shape a phytoplankton assembly and how phytoplankton will respond or evolve to future ocean environments in the context of climate change in which P supply from deep ocean is predicted to decrease. The mutants generated here will be a valuable resource for future studies to further dissecting the molecular machinery underlying phytoplankton acclimation and adaptation to P variability (5, 37).

**Fig. 4.**
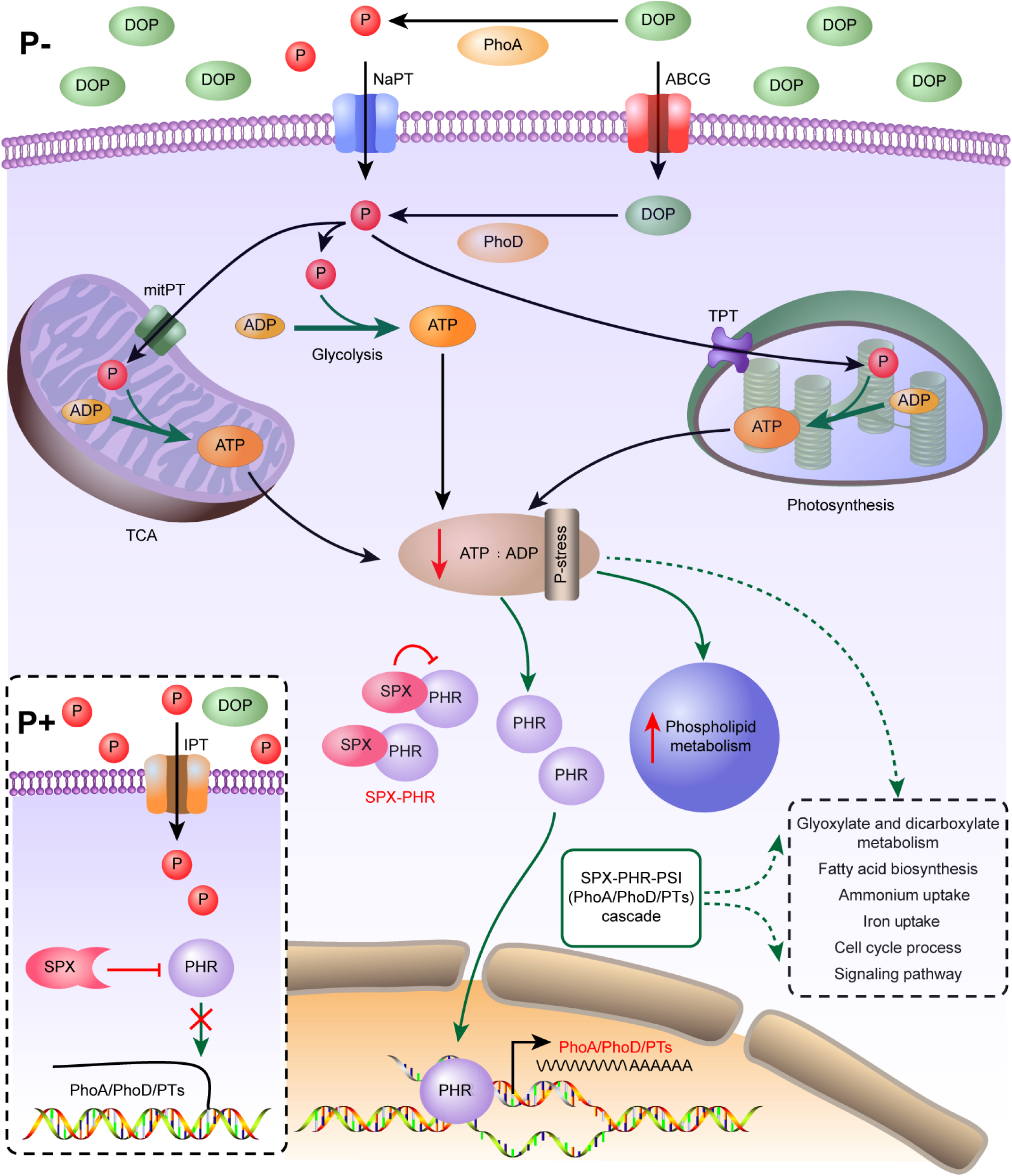
Draft P metabolic atlas in the model diatom under P+ and P− conditions. Underlined and italicized are various metabolic processes. The P+ condition is shown in the inset (lower left), while the main figure depicts P− condition. Positive and negative regulations are indicated by arrows and flat-ended lines, respectively. Vertical red arrows indicate up- and down-regulations, respectively.

### SPX-PHR regulatory cascade of AP and DIP transporter expression

Our present results and the literature converge at a central P regulatory cascade (Fig. 4), which simplistically can be presented as SPX-PHR-PSI, where PSI (P starvation induced) represents AP and PT as well as P-responsive effectors not discussed in this paper, SPX is an inhibitory master regulator, and PHR is a secondary regulator that mediates regulating signals of SPX to modulate expression of PSI. These regulatory effects deeply influence many metabolic pathways through P-dependent or AP-dependent control circuits.

The SPX protein is known in plants as a regulator of P homeostasis and P signalling (38, 39). Its role in *P. tricornutum* as a master regulator of P acquisition and homeostasis is demonstrated by multiple lines of evidence (Fig. 4). First, SPX mutation leads to significant up-regulation of AP and PT genes, even under P+ conditions that would usually down-regulate these genes, indicating that SPX is an inhibitory up-stream regulator. Second, *SPX* transcription is responsive to P stress, as expected of a P regulator that senses internal P levels and controls its uptake (40, 41). Finally, WGCNA analysis reveals the dominant and broad metabolic influences of SPX.

In plants, SPX functions through PHR, a positive regulator in P signaling (24, 40, 42). PHRs are transcription factors, harbouring myeloblastosis (MYB) and coiled-coil (CC) domains for DNA binding (43, 44). The response of the homologs in *P. tricornutum* WT to P stress and its lack of response in *m*PhoA and *m*PhoD places PHR as an intermediate between SPX and effectors. Furthermore, as recently reported, its knockout results in a significant decline of AP expression, demonstrating that PHR is a positive regulator (25). It is plausible then that under P+ conditions, SPX depresses PHR, thus maintaining low expression of AP and PT (Fig. 4). Under P stress, we find a greater increase in *PHR* than *SPX* expression, causing the PHR:SPX transcript ratio to increase from 1:9 under P+ to 1:4 under P− conditions (Table S18). As SPX presumably functions through binding to PHR (24), the relative increase of PHR may weaken the inhibitory signal of SPX, leading to elevated AP and PT expression (Fig. 4). This explains the observed counterintuitive up-regulation of SPX under P− conditions. To confirm this hypothesis, further study is warranted.

### Functional differentiation and compensatory regulation of APs and PTs confers versatility and flexibility in scavenging DOP

Our comparative transcriptomic analyses reveal P-responsive and non-P-responsive APs, the latter likely functioning in regular metabolic pathways (Table S4). Similarly, PTs also diverge into high-affinity or low-affinity groups (Fig. S7). Among the P-responsive APs, our data clearly show that PhoA is an extracellular AP, verifying a previous report (35), usually accounting for ∼75% of total AP activity, whereas PhoD is intracellular, accounting for <25%. More importantly, PhoA and PhoD show functional propensities towards different types of DOP substrates: when *P. tricornutum* is provided G3P, PhoA hydrolyses G3P extracellularly to release PO43-for uptake, but when phytate is the sole P source, it is imported for the intracellular PhoD to harvest. Such functional specialization between APs has previously not been documented, and hence should be investigated more broadly across phytoplankton phylogenetically tree. It is noteworthy that *P. tricornutum* utilizes phytate via AP, while other organisms do so via phytase (45, 46); our search of the *P. tricornutum* genome returned no phytase gene.

Our data further show that the functional differentiation between PhoA and PhoD is not static but strikingly dynamic. Although PhoA is usually responsible for G3P utilization, when it is functionally lost, the usually minor PhoD is transcriptionally up-regulated to maintain G3P utilization. Similarly, the disruption of *PhoD* induces elevation of *PhoA* expression. This is an intriguing combination of functional replacement and compensatory up-regulation of one AP for another.

These characteristics have significant ecological and evolutionary implications (Fig. 1e). First, the functional differentiation and the ability to switch APs give the alga the flexibility to utilize different types of DOP available in the environment. Notably, utilization of phytate does not yield the same flexibility as G3P; when the primary responsible enzyme PhoD is lost, PhoA does not step up to enable phytate utilization. Second, the bidirectional compensatory expression scheme allows the phytoplankton to modulate expression levels of different APs for utilizing different concentrations and types of available DOPs. Conceivably, when PhoA is switched to PhoD as in the case of *PhoA* mutation, a transporter of the DOP substrate is needed. Hence, the higher efficiency of utilizing DOP is off set by the cost in synthesizing DOP transporter in the case of PhoD, and the benefit of hydrolysing DOP extracellularly without having to synthesise transporters is off set by losing part of the resulting PO43-in the case of PhoA. It follows that some DOPs may be too costly to import, giving advantages to extracellular hydrolysis. Third, some DOPs such as TEP, not accessible to *P. tricornutum*, pose additional factors influencing phytoplankton community assembly by favoring species able to utilize these DOPs. Finally, the ability to switch between different APs for using one DOP substrate may relax constraints for AP mutation, possibly the reason that AP sequences rapidly diverged (5).

### Metabolic landscape influenced by APs and P and potential P-dependent iron uptake

The combination of gene knockouts and transcriptome profiling unveils remarkable influences of APs, SPX, and P conditions on the metabolic landscape (Fig. 4). This is more deeply demonstrated by our WGCNA analyses, which help disentangle compounded effects of P condition, AP, and other SPX-dependent factors. Data (Table S19) show that the SPX-PHR-AP cascade influences glyoxylate/dicarboxylate metabolism (involving PhoA) and ammonium uptake/arginine metabolism (involving non-PhoD); fatty acid biosynthesis is positively influenced by P deficiency while long-chain lipid synthesis is negatively regulated by PhoA; phospholipid metabolism is under the influence of PhoD-PhoA balance and AP-PT regulation; photosynthesis is regulated by P condition and some APs; glycolysis and TCA cycle are influenced by P stress or APs under the control of SPX at different nodes of the pathway; and SPX and non-PhoA, non-PhoD APs may be involved in regulating iron uptake under P-deficiency. These indicate that APs have dual functions, one in scavenging DOPs and the other in regulating metabolic processes under normal P conditions. It will be fruitful to further dissect the influences of AP and P condition on these metabolic pathways, particularly iron uptake, photosynthetic carbon fixation, and the cell cycle (governing intrinsic population growth) in phytoplankton.

## Data availability

Raw sequencing data is now available at NCBI in the SRA (Short Read Archive) database under accession number SRP214503.

## Author contribution

S. Lin conceived and supervised the work. Zhang, Zhou, S. Lin, Li, and X. Lin designed the experiments and data analysis strategies. Zhang, Wang, Li, Wu, and You carried out the experiments. Zhang and Zhou conducted data analyses. Zhang and S. Lin wrote the manuscript. All authors participated in revising the manuscript and agreed to the final submitted version.

## Acknowledgements

We are indebted to Drs. Chris Bowler (IBENS, France) and Jacquine Niles (MIT, USA) for sharing experience in and providing suggestions on gene transformation or CRISPR/Cas9. We also thank Dr. Xin Lin (Xiao), a diatom epigenetics expert, for providing the plasmids and Ms. Chentao Guo for her logistic support. The work was financially supported by the Marine S & T Fund of Shandong Province for Pilot National Laboratory for Marine Science and Technology (Qingdao) (grant # 2018SDKJ0406-3) and Natural Science Foundation of China (grant #41776116). The Marine Microbial Initiative (MMI) of the Gordon and Betty Moore Foundation provided funding toward developing protist functional tool (grant #GBMF grant #4980.01) that enabled S. Lin to develop this work.

