## Supplementary figures and tables for "One enzyme many faces: alkaline phosphatase-based phosphorus-nutrient strategies and the regulatory cascade revealed by CRISPR/Cas9 gene knockout": Supplementary Figures.pdf

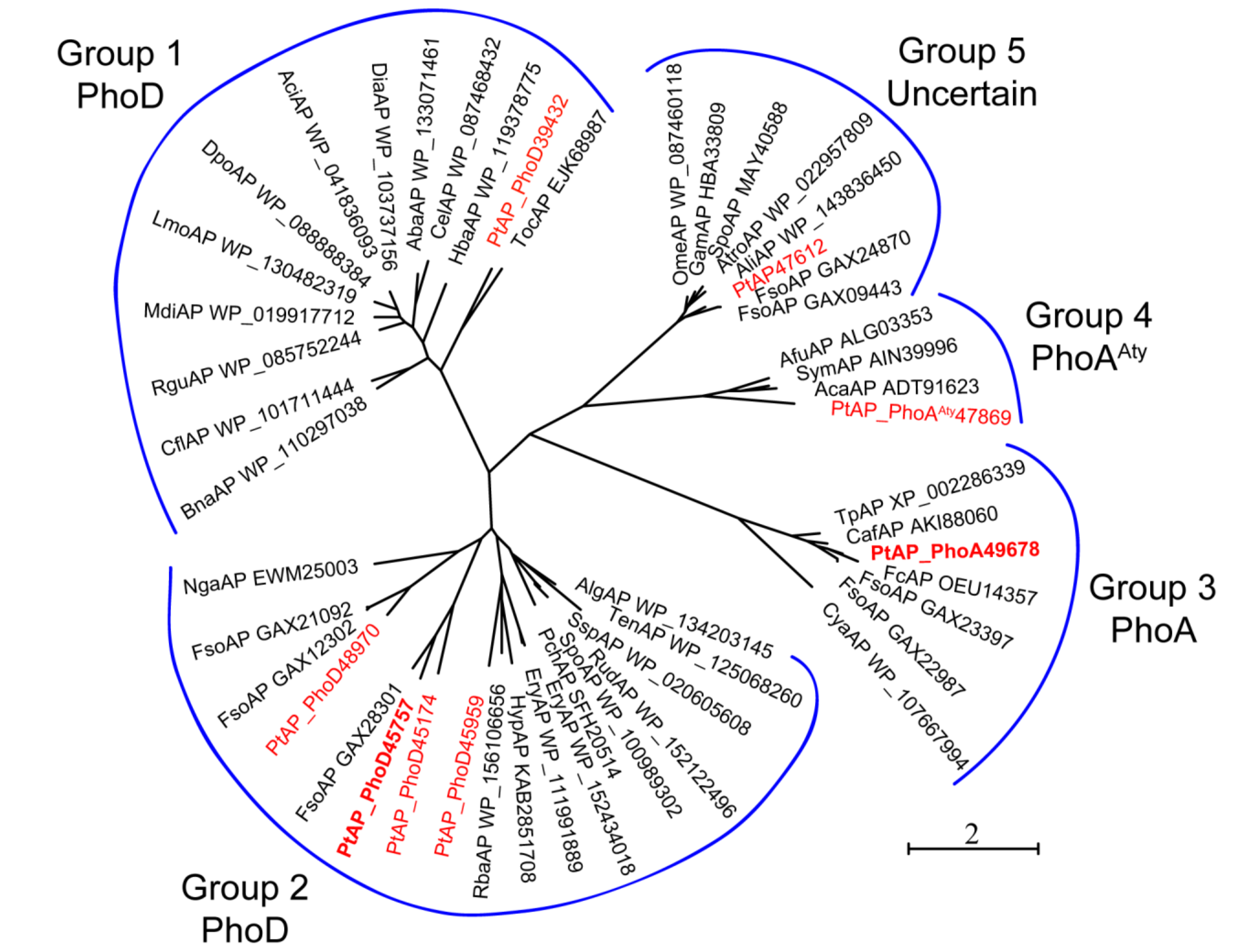


**Figure S1. Phylogenetic relationship of AP genes identified from *P. tricornutum* with known types of APs.** The eight *P. tricornutum* AP genes were assigned to 5 groups including one PhoA, five putative PhoD, one atypical PhoA and one unclassified AP genes. Scale bar depicts substitution rates per amino acid residue. Group 1 (PhoD) contains *Thalassiosira oceanica* (TocAP EJK68987), *P. tricornutum* (PtAP39432), *Henriciella barbarensis* (HbaAP WP_119378775), *Cellvibrio sp. PSBB006* (CelAP WP_087468432), *Alteromonadaceae bacterium M269* (AbaAP WP_133071461), *Diaphorobacter sp. LR2014-1* (DiaAP WP_103737156), *Acidovorax sp. JS42* (AciAP WP_041836093), *Diaphorobacter polyhydroxybutyrativorans* (DpoAP WP_088888384), *Leptothrix mobilis* (LmoAP WP_130482319), *Methyloversatilis discipulorum* (MdiAP WP_019917712), *Rhizobacter gummiphilus* (RguAP WP_085752244), *Caulobacter flavus* (CflAP WP_101711444), *Blastomonas natatorial* (BnaAP WP_110297038). The Group 2 (PhoD) contains *Nannochloropsis gaditana* (NgaAP EWM25003), *Fistulifera solaris* (FsoAP GAX21092), *F. solaris* (FsoAP GAX12302), *P. tricornutum* (PtAP48970), *F. solaris* (FsoAP GAX28301), *P. tricornutum* (PtAP45757), *P. tricornutum* (PtAP45174), *P. tricornutum* (PtAP45959), *Rhizobiales bacterium YIM 77505* (RbaAP WP_156106656), *Hyphomicrobiaceae bacterium* (HypAP KAB2851708), *Erythrobacter sp. KY5* (EryAP WP_111991889), *Erythrobacter sp. THAF29* (EryAP WP_152434018), *Pontibacter chinhatensis* (PchAP SFH20514), *Spirosoma pollinicola* (SpoAP WP_100989302), *Rudanella sp. HX-22-17* (RudAP WP_152122496), *Spirosoma spitsbergense* (SspAP WP_020605608), *Tenacibaculum sp. DSM 106434* (TenAP WP_125068260), *Algoriphagus sp. ARW1R1* (AlgAP WP_134203145). The Group 3 (PhoA) contains *P. tricornutum* (PtAP49678), *Thalassiosira pseudonana* (TpAP XP_002286339), *Chaetoceros affinis* (CafAP AKI88060), *Fragilariopsis cylindrus CCMP1102* (FcAP OEU14357), *F. solaris* (FsoAP GAX23397), *F. solaris* (FsoAP GAX22987), *Cyanothece sp. BG0011* (CyaAP WP_107667994). The Group 4 (PhoA$^{Aty}$) contains *Alexandrium fundyense* (AfuAP ALG03353), *Symbiodinium sp. CCMA192* (SymAP AIN39996); *Amphidinium carterae* (AcaAP ADT91623), *P. tricornutum* (PtAP47869). The Group 5 (unclassified) contains *Oleiphilus messinensis* (OmeAP WP_087460118), *Gammaproteobacteria* (GamAP HBA33809), *Spongiibacter sp.* (SpoAP MAY40588), *Spongiibacter tropicus* (AtroAP WP_022957809), *Aliiglaciecola sp. M165* (AliAP WP_143836450), *F. solaris* (FsoAP GAX24870), *F. solaris* (FsoAP GAX09443), *P. tricornutum* (PtAP47612).

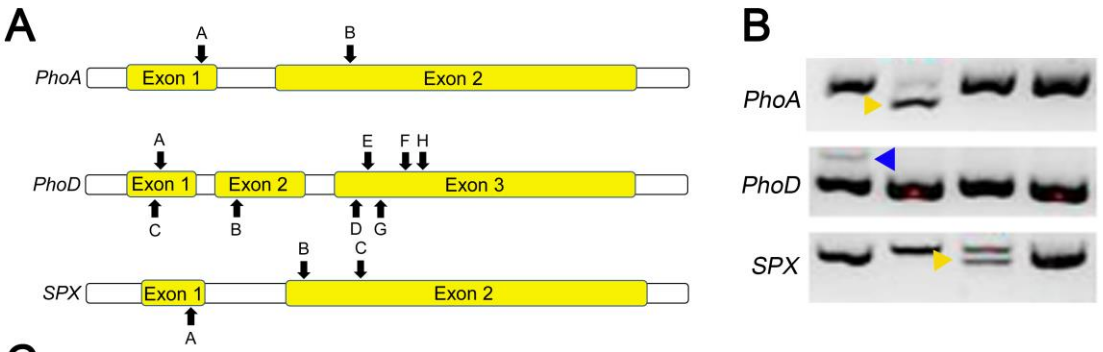


C

| Gene | Targeting sgRNAs | Clone number | Targeted number | Targeting efficiency (%) | Mediated repairs |
|---|---|---|---|---|---|
| *PhoA* | sgRNA2 | 59 | 25 | 42.4 | NHEJ |
| *PhoD* | sgRNA8 | 88 | 30 | 34.1 | NHEJ |
| *SPX* | sgRNA3 | 54 | 5 | 9.3 | NHEJ |

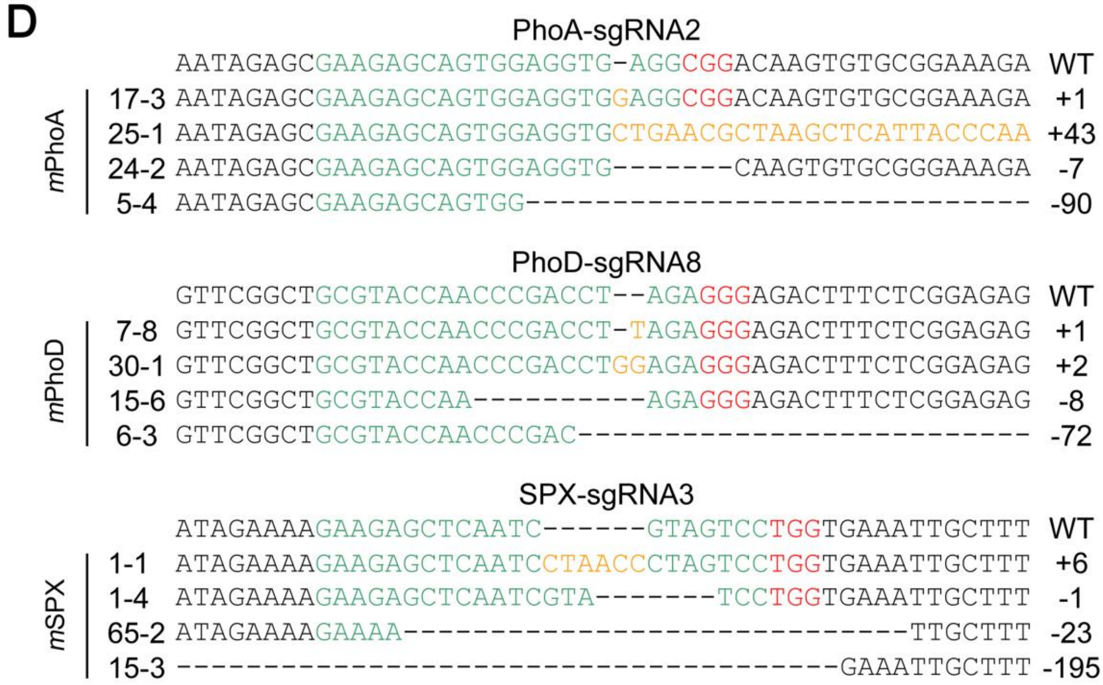


**Figure S2. Successful knockout of two AP genes (*PhoA* and *PhoD*) and the gene of SPX protein. (A)** Schematic diagram of sgRNA-targeting sites. Letters above bar (A to H) represent the targeting sites from sgRNA1 to sgRNA8, respectively. **(B)** Target regions of these three genes amplified with specific primers, with the yellow arrowheads showing deletion mutations and the blue arrowhead showing insertion mutation. **(C)** Summary of on-target rates (targeting efficiencies) in *m*PhoA, *m*PhoD and *m*SPX. NHEJ，non-homologous end joining. **(D)** Representative sequencing results of mutant clones in the PhoA locus using sgRNA2, in the PhoD locus using sgRNA8 and in the SPX locus using sgRNA3, respectively. The PAM sequences and inserted sequences are labeled in red and yellow, respectively.

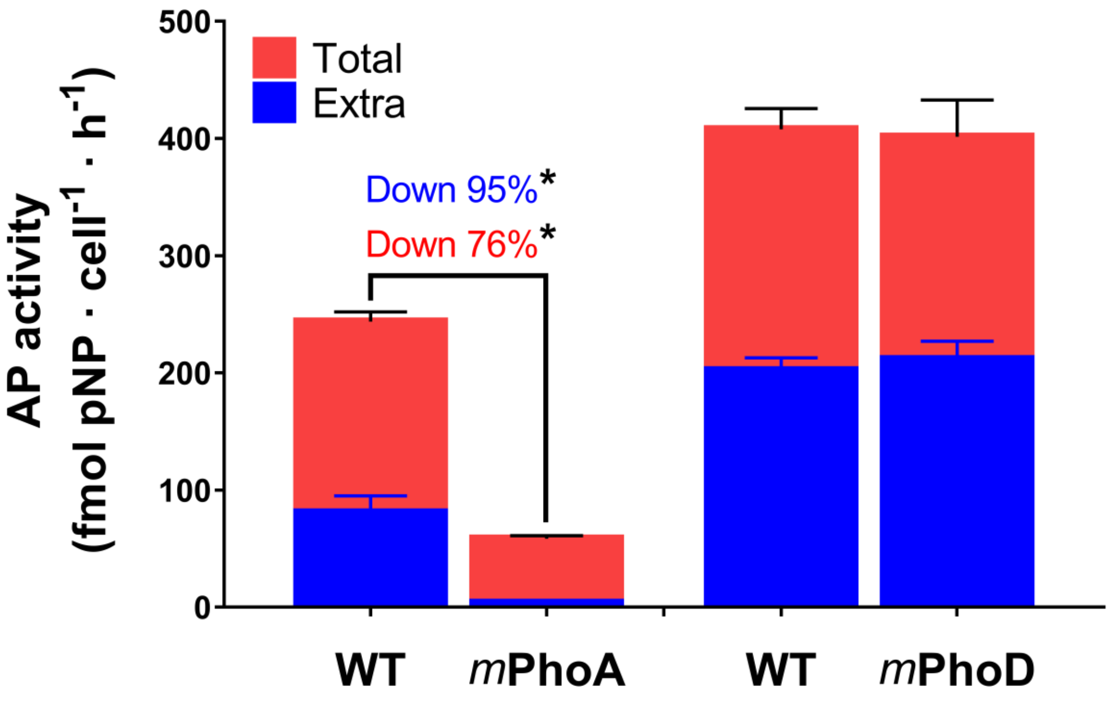


**Figure S3. Changes of alkaline phosphatase activities after mutation of PhoA and PhoD.** Data are from day 7 of the P-deprived *m*PhoA and *m*PhoD cultures. The total (tAP, red bar) and extracellular (exAP, blue bar) AP activities decreased significantly after PhoA mutation (compared with wild-type or WT). However, PhoD mutation caused no significant change in AP activity in *m*PhoD. Asterisk shows significant difference.

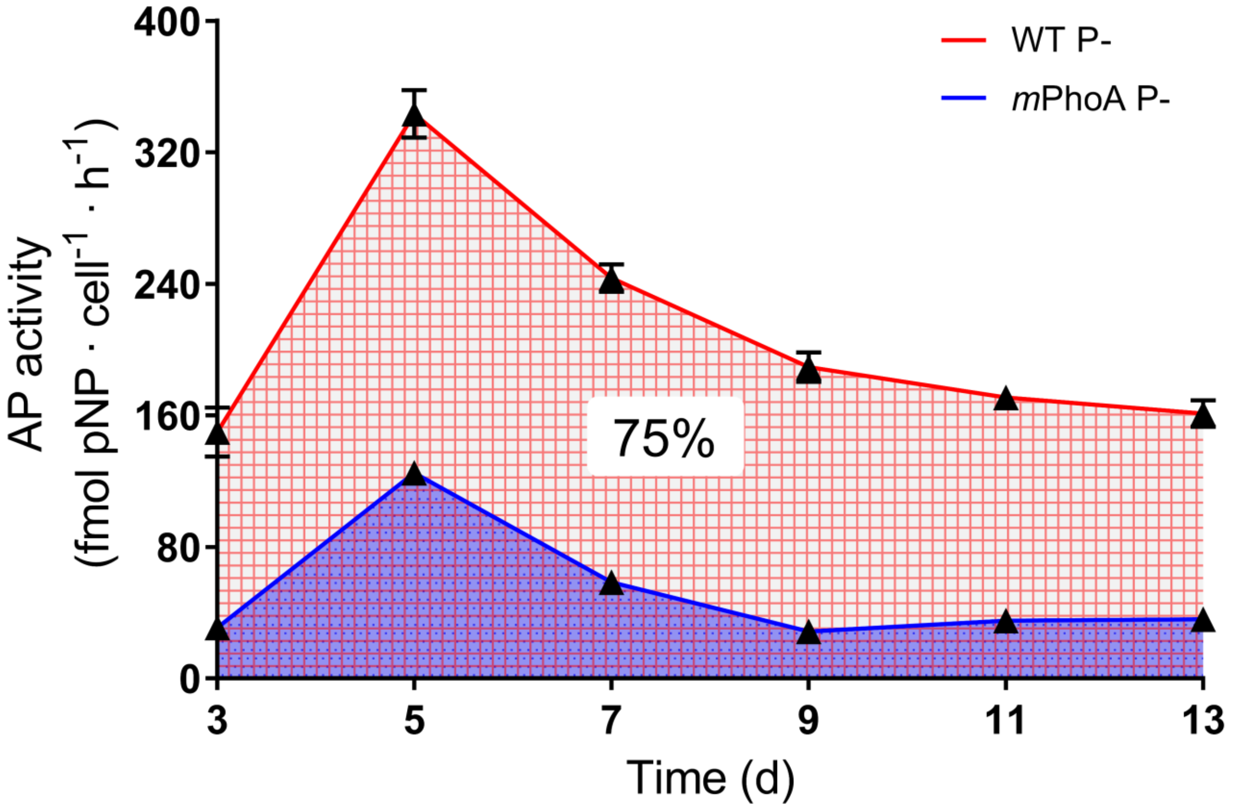


**Figure S4. Temporal trends of total alkaline phosphatase activities in wild-type (WT) and PhoA mutant (*m*PhoA) grown under P-stress (P−) condition**. When averaged over the 10-day experimental period, PhoA accounted for ~75% of the total AP activity. Each data point is the mean of triplicate cultures with the error bar indicating the standard deviation.

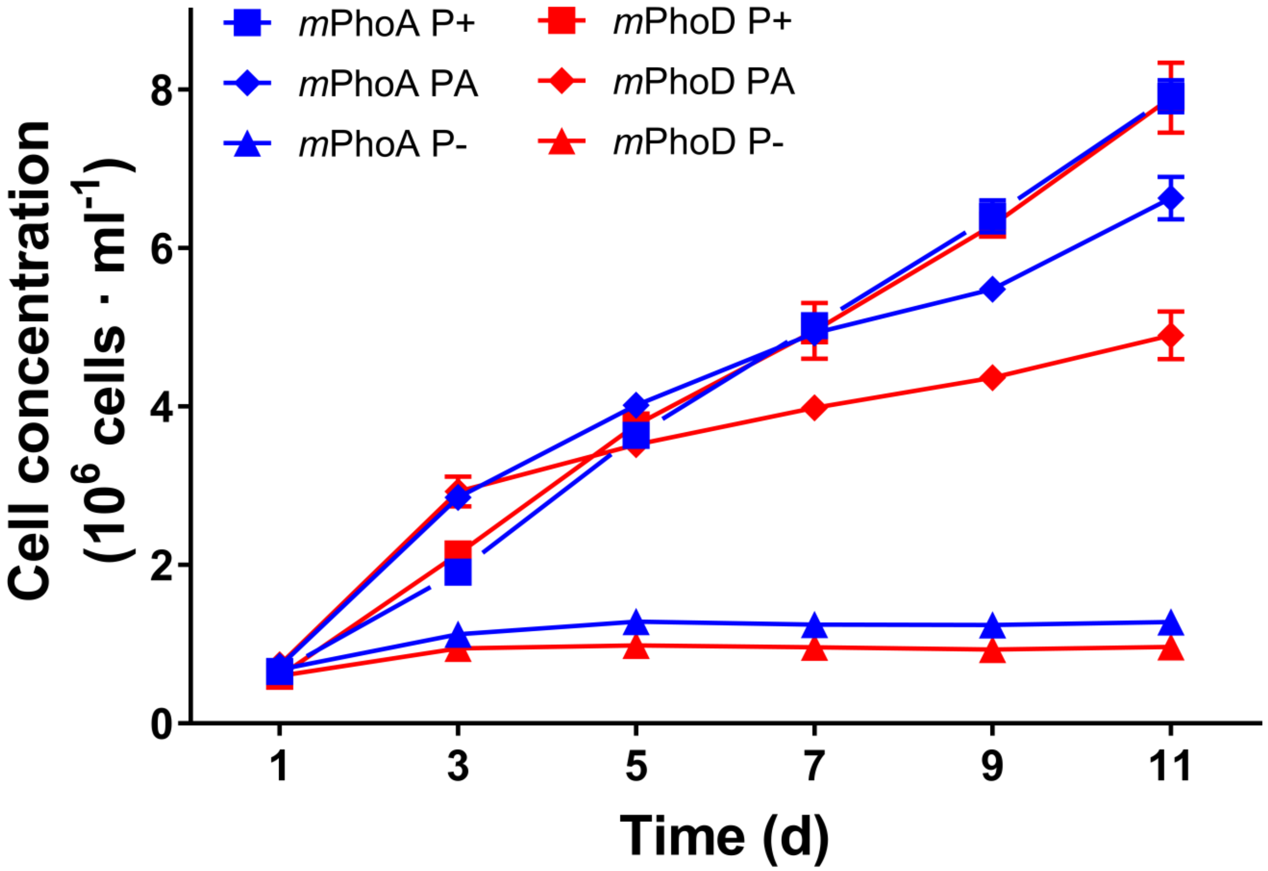


**Figure S5. Growth curves of PhoA and PhoD mutant (*m*PhoA and *m*PhoD) strains under different P conditions.** P+, P-replete; P−, P-depleted; PA, phytate. Each data point is the mean of triplicate cultures with the error bar indicating the standard deviation.

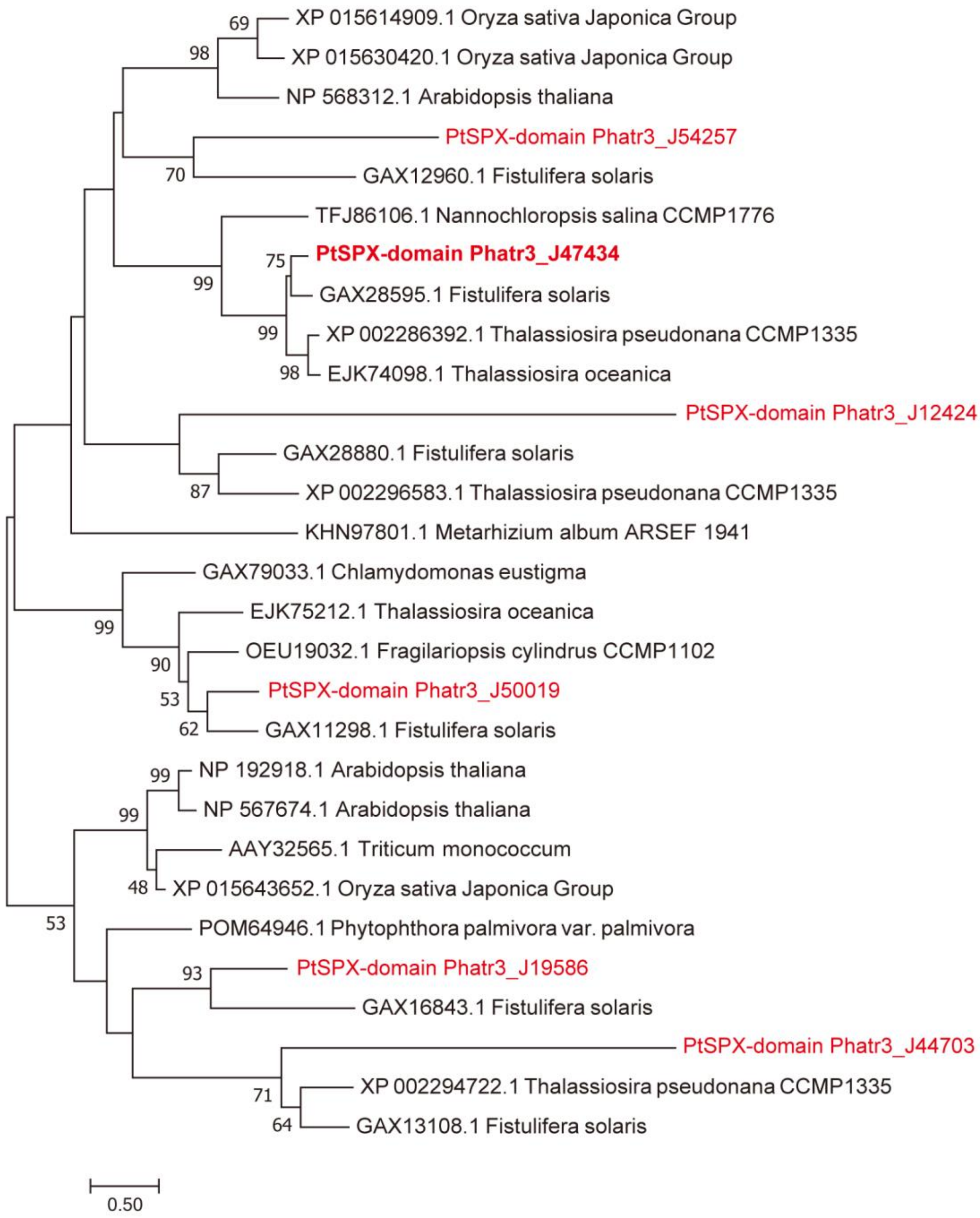


**Figure S6. Maximum likelihood tree based on amino acid sequences of SPX domains previously reported in plants (*Arabidopsis*, Oryza, *Triticum*) and those identified in this study from algae.** The SPX-domain sequences in *P. tricornutum a*re marked in red and the bold-typed one (Phatr3_J47434) was mutated in our study. Values at nodes were bootstrap support values (only those > 50% are shown). Scale bar depicts substitution rate per amino acid residue.

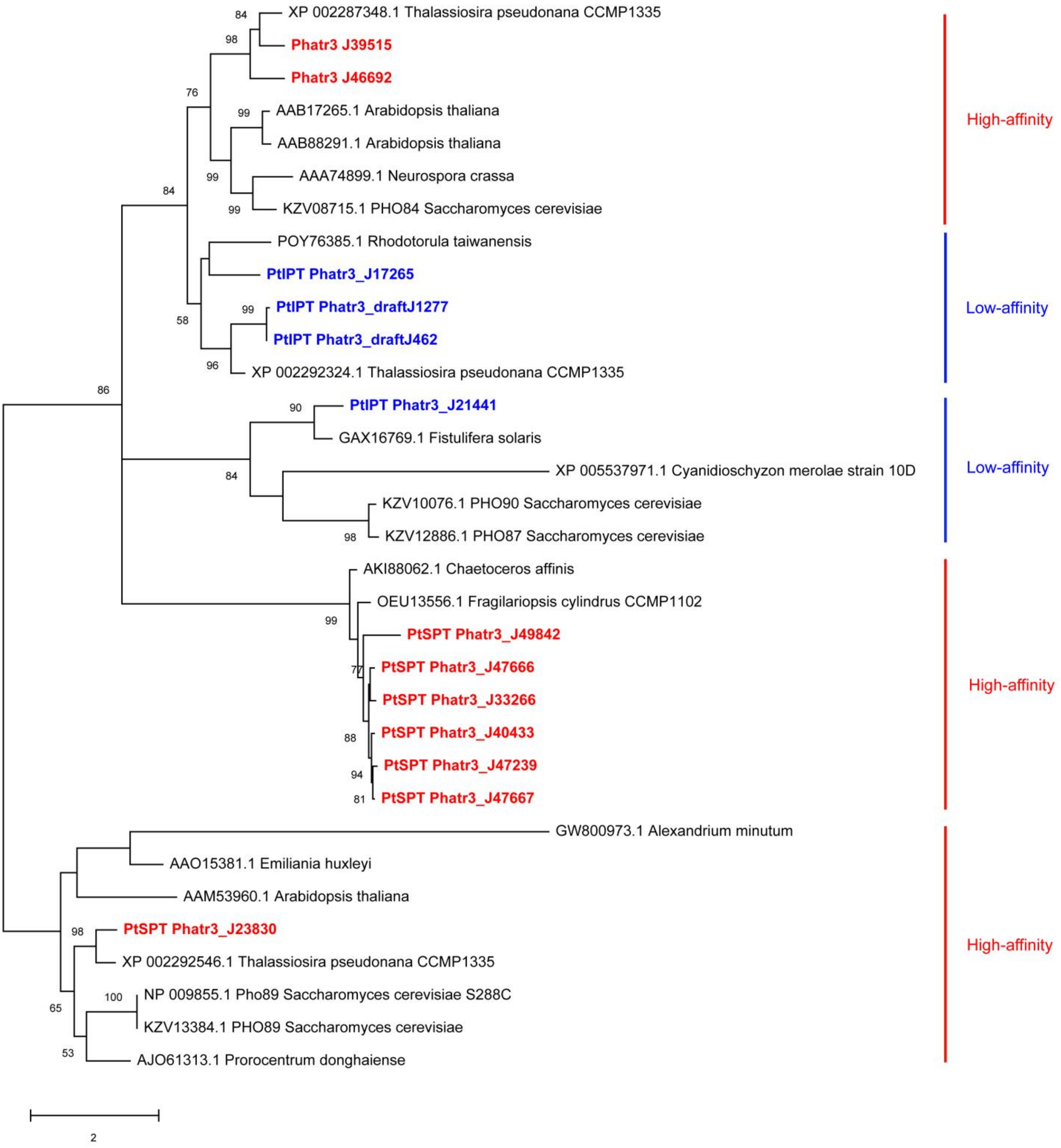


**Figure S7. Phylogenetic relationship of phosphate transporter (PT) genes identified in *P. tricornutum* with known types of PTs.** Nine of the 13 *P. tricornutum* PT genes were clustered with high-affinity transporters (in red) and the other four to low-affinity transporters (in blue). Notably, all the seven sodium phosphate co-transporters (SPTs) are of high affinity and most (four) of the six inorganic phosphate transporters (IPTs) are of low affinity. Values at nodes were bootstrap support values (only those > 50% are shown). Scale bar depicts substitution rate per amino acid residue.

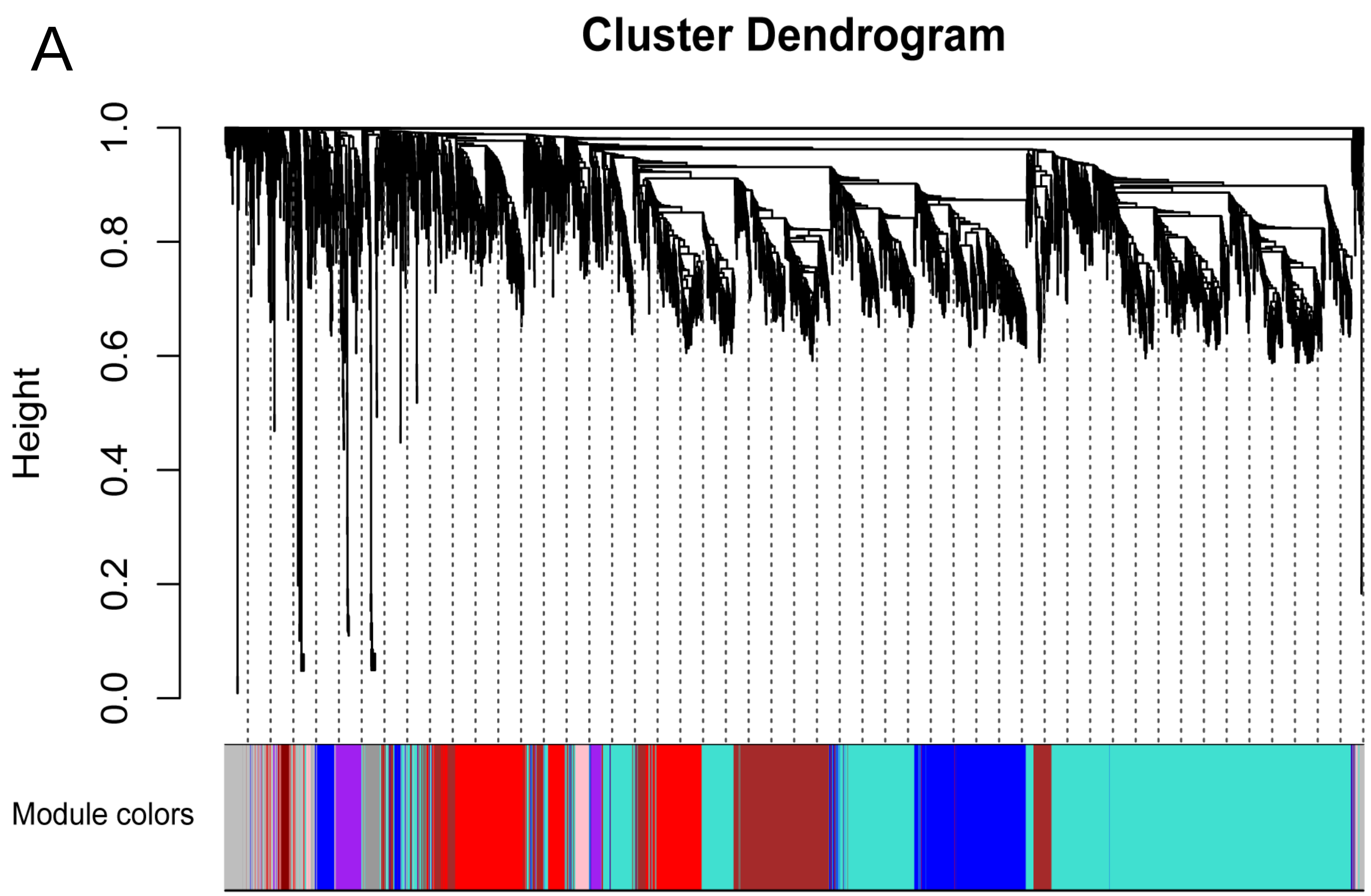


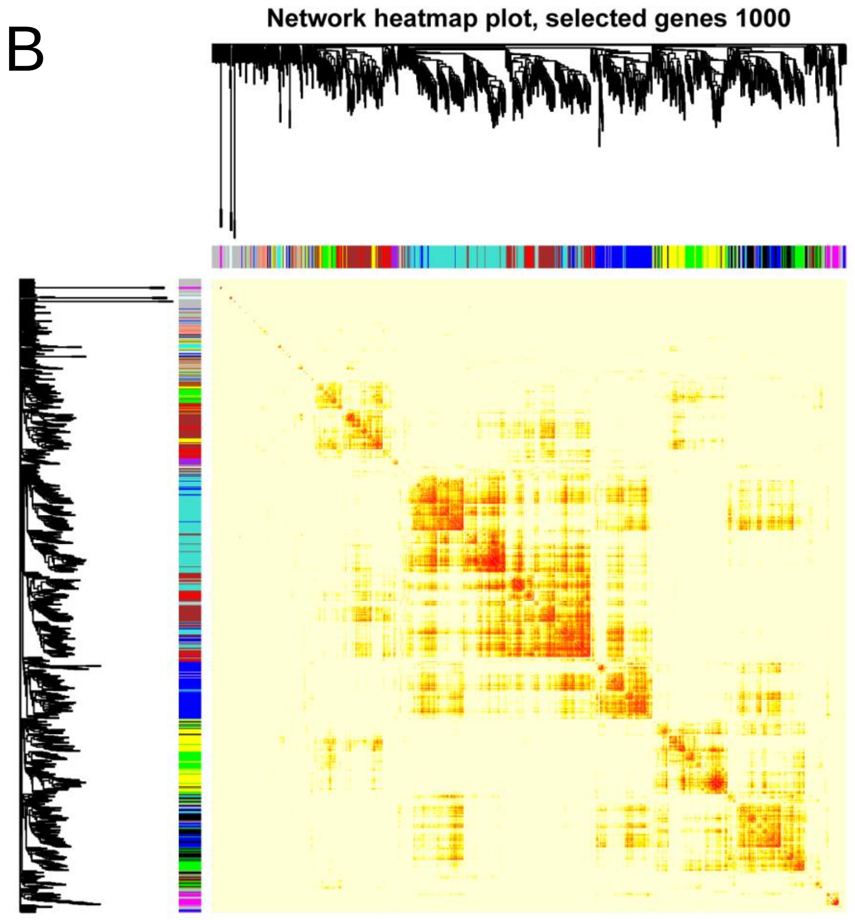


**Figure S8. Weighted gene co-expression network analysis (WGCNA) based on AP activities.** **(A)** Gene dendrogram obtained by average linkage hierarchical clustering. Gene expression similarity is determined using a pair-wise weighted correlation metric, and clustered according to a topological overlap metric into modules; assigned modules are colored on bottom. **(B)** Network heatmap plot. Each row and column corresponds to a gene, and high co-expression interconnectedness is indicated by progressively more saturated yellow and red colors. The gene dendrogram and module assignment are shown on the left and top.
